# Filopodia numbers impact chemotactic migration speed

**DOI:** 10.1101/2025.05.28.656686

**Authors:** Annika L. Schroder, Livia D. Songster, Yahor Savich, Margaret A Titus

## Abstract

Migrating cells sense and respond to external chemical and physical cues, enabling them to efficiently reach their destinations. Filopodia are slender actin-filled membrane protrusions implicated in interacting with the extracellular environment in many contexts, such as neuronal growth cone guidance and the capture of prey by immune cells and unicellular organisms. The role of filopodia in chemotactic guidance in fast-moving amoeboid cells has not been well-studied. The social amoeba *Dictyostelium* relies on chemotaxis for development and finding food, making it an excellent system for investigating the role of filopodia in amoeboid chemotaxis. Stimulation of amoebae with the chemoattractant cAMP activates a transient increase of filopodia formation by recruiting the filopodial myosin DdMyo7 to the cell cortex. Filopodia formation is biased towards the source of chemoattractant, yet *myo7* null cells that lack filopodia exhibit normal directional migration. However, cells either lacking filopodia or having increased numbers of filopodia move more slowly than those with wildtype numbers of filopodia. Thus, while filopodia are dispensable for detection of chemical gradients by amoeboid cells, changes in filopodia number can impact their migration speed possibly due to altering cell-substrate adhesion.

**In brief:** Filopodia formation in chemotactic amoeboid cells is stimulated by chemoattractant and biased towards the gradient source. Lack of filopodia does not impair directed migration but rather reduces cell speed. Changes in filopodia number correlate with the speed of chemotactic cells suggesting a role for these extension in tuning adhesion for optimal migration.

**Highlights:**

- Filopodia formation in chemotactic amoeboid cells is biased towards the source of the gradient.
- Chemotactic *Dictyostelium* amoebae lacking filopodia migrate with directional persistence.
- Migration speed is altered by filopodia number.

## Introduction

Cells navigate diverse and complex environments when seeking food, repairing wounds or coming together to form multicellular structures such as tissues or neural networks. Their ability to move efficiently through the environment to their target relies on the cell’s ability to detect both chemical and physical cues. Filopodia are slender membrane protrusions supported by parallel bundles of actin filaments that serve as cellular sensors (Heckman and Plummer, 2013). Diverse cell types make filopodia, including neurons, epithelial cells, endothelial cells (Gallop, 2020) and unicellular organisms such as *Capsaspora*, choanoflagellates such as *Salpingoeca rosetta* (Sebé-Pedrós et al., 2013) and *Dictyostelium* amoebae (Kobilinsky et al., 1976). The role of filopodia in chemotactic guidance has been widely studied in mammalian systems, most notably in neurons where early work revealed that their loss results in disorientation of neural growth cones during embryonic development (Bentley and Toroian-Raymond, 1986, Chien et al., 1993). Filopodia have also been implicated in the directional migration of endothelial cells towards BMP (Pi et al., 2007). Filopodia can have a role in the interaction of cells with the extracellular matrix to reorganize fibronectin, such as during somite morphogenesis in quail embryogenesis (Sato et al., 2017) or migration of collectives of cancer cells (Summerbell et al., 2020) or sensing substrate stiffness (Wong et al., 2014). While much is known about filopodia function in mesenchymal cells, the role of these projections in fast-moving chemotactic amoeboid cells is not well-characterized.

The social amoeba *Dictyostelium* is an excellent system for investigating the function of filopodia in amoeboid migration and chemotaxis. The vegetative amoebae move randomly, producing filopodia that extend from dynamically protruding actin-rich regions of the cell (Tuxworth et al., 2001, Petersen et al., 2016). The MyTH-FERM (<u>my</u>osin tail <u>h</u>omology 4 - band <u>4</u>.1 <u>e</u>zrin <u>r</u>adixin <u>m</u>oesin) DdMyo7 and the actin polymerase VASP are both strongly localized to the tips of filopodia and loss of either results in a significant reduction in filopodia formation in vegetative *Dictyostelium*, as well as a decrease in substrate adhesion and phagocytosis (Tuxworth et al., 2001, Han et al., 2002). Upon starvation, *Dictyostelium* execute a developmental program that promotes the aggregation of cells into a mound by directed migration to the secreted chemoattractant cAMP. The chemotactic signaling pathways and cytoskeletal changes essential for this process are well characterized (Artemenko et al., 2014, Pal et al., 2019) yet it is not known if these pathways are linked to filopodia formation and if filopodia have a role in chemotactic migration in *Dictyostelium* or other fast-moving amoeboid cells. The inability of the *myo7* and *vasp* null mutants to make filopodia makes them well-suited to investigate the contribution of these special structures to the migration and chemotactic efficiency amoeboid cells.

## Results & Discussion

The adhesion defect of *myo7* mutants causes these cells to frequently lose contact with the substrate during migration (Tuxworth et al., 2001), potentially resulting in changes in the cell centroid that might confound measurement of speed. Migration and chemotaxis were thus measured using an under-agarose assay (Laevsky and Knecht, 2001), using 0.7 - 0.8% agarose that kept the cells confined but did not compress them. Cells moving under this percentage of agarose make normal numbers of filopodia and do not bleb (Tyson et al., 2014). The migration of control Ax3 cells in both nutrient media (HL5) and phosphate starvation buffer (PB) used for when analyzing the speed of chemotactic cells was first measured. There are differences in cell speed in the two different conditions: control cells under agarose move significantly slower in HL5 than in in PB (4.0 µm/min vs 11.5 µm/min; Fig 1), suggesting that components in the rich growth medium alter the interaction of cells with the substrate and reduce their speed. The *myo7* null mutant lacking filopodia moved at the same speed as control cells in HL5, but at half the speed of control cells in PB (Fig 1). Interestingly, *myo7* null cells expressing a mutant DdMyo7 defective in autoinhibition that results in increased filopodia formation (KKAA; cells make a 2-fold excess of filopodia (Petersen et al., 2016, Arthur et al., 2019)) also move at the same speed as control cells in HL5 but at half the speed of control cells in PB. These results reveal that filopodia play a role in migration and that both too many and too few filopodia reduce the rate of cell speed.

**Figure 1.**
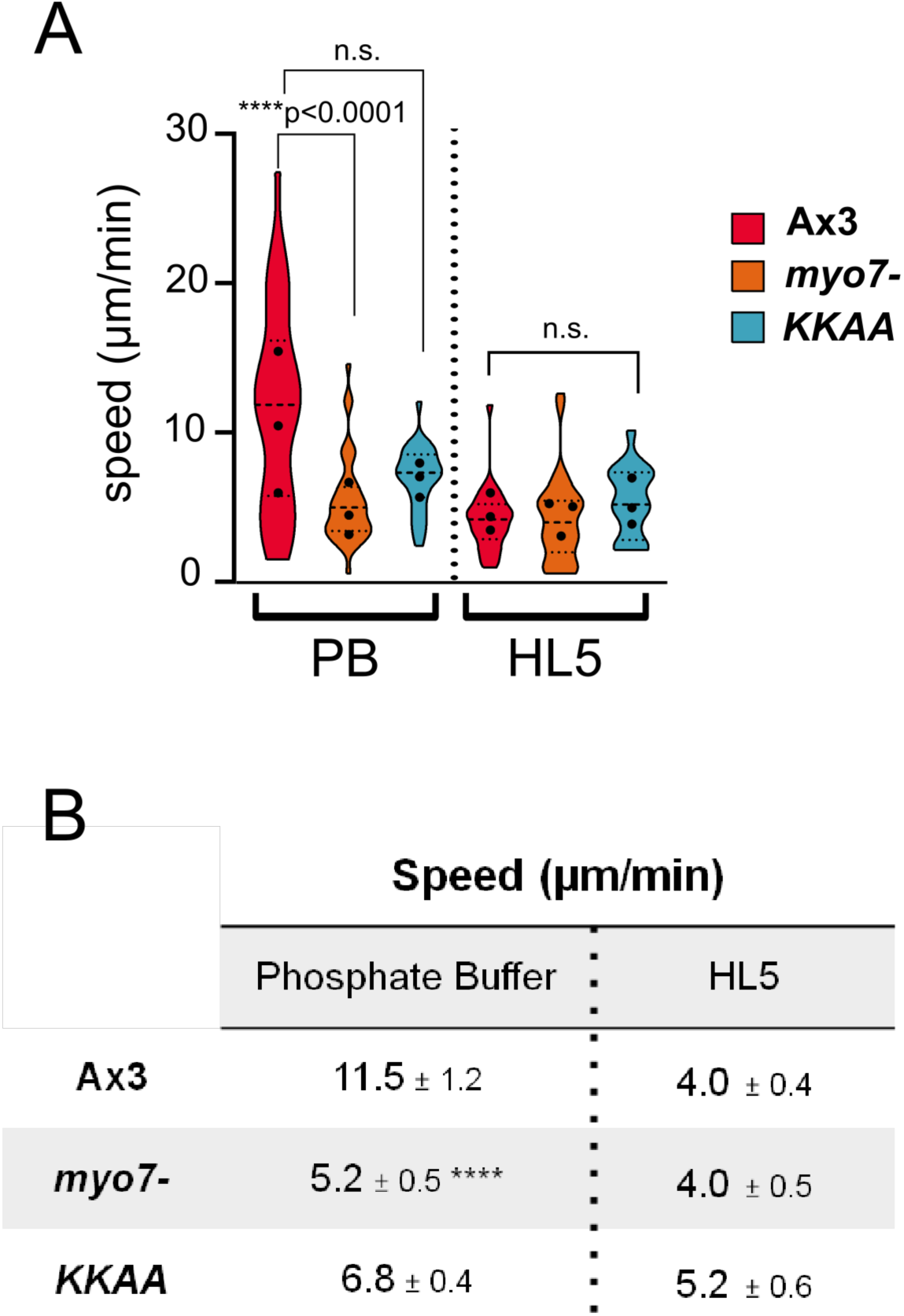
Filopodia number impact the speed of randomly moving cells. **A.** Average speed of vegetative wild type (Ax3), *myo7* null or *myo7* nulls expressing the KKAA autoinhibition mutant when assayed under agarose. Shown is the speed of cells when placed in phosphate starvation buffer (PB) or in nutrient media (HL5). N= 3, n = 16 (minimum) for each assay. Kruskal-Wallis ANOVA and multiple comparisons tests (Ax3: ****p<0.0001, n.s.= not significant, p=0.067; Ax2, n.s., p=0.14) **B.** Summary of means values and significance measured for each line ± SEM.)

### Chemoattractant stimulation triggers filopodia production

Global stimulation of chemotactically competent *Dictyostelium* with the chemoattractant cAMP evokes a characteristic sequence of signaling and cytoskeletal changes that first results in increased cortical levels of F-actin accompanied by cellular contraction (5 sec), followed by random pseudopod extension (35 sec) (McRobbie and Newell, 1983, Hall et al., 1988). Stimulation of wild type cells expressing GFP-DdMyo7 (hereafter DdMyo7) results in rapid recruitment of the filopodial myosin to the cell cortex, with a clear single peak occurring at 4.5 sec (Fig. 2A; Supp Fig 1). DdMyo7 recruitment was concurrent with that of PIP3 as visualized by GFP-CRAC (<u>c</u>ytosolic <u>r</u>egulator of <u>a</u>denylyl <u>c</u>yclase), and slightly preceded the robust initial peak of actin recruitment (Fig. 2A-B) (Parent et al., 1998, Sasaki et al., 2004). The second, smaller peak of cortical actin is not accompanied by an increase in cortical DdMyo7. mApple-VASP is also recruited to the cortex following global stimulation with cAMP, with a single peak at ∼ 4 sec, similar to what is observed for DdMyo7 (Supp Fig 2A, B). Treatment of cells with Latrunculin A (latA) prior to cAMP stimulation abolished cortical recruitment of both DdMyo7 and mApple-VASP (Supp Fig. 2C-D) establishing that actin is required for recruitment of both key filopodial proteins. The dramatic recruitment of DdMyo7 and VASP to the cortex was accompanied by a transient increase in filopodia number, with a peak ∼1.5-fold increase at 5.8 sec post stimulation (Fig. 2C). The cAMP-stimulated filopodia appear to extend out of local patches of cortical GFP-DdMyo7 (Fig. 2A). Thus, chemotactic stimulation results in the rapid recruitment of key filopodial proteins to the cortex, promoting increased filopodia production (Fig. 2D) and suggesting that filopodia may play role in directed cell migration.

**Figure 2.**
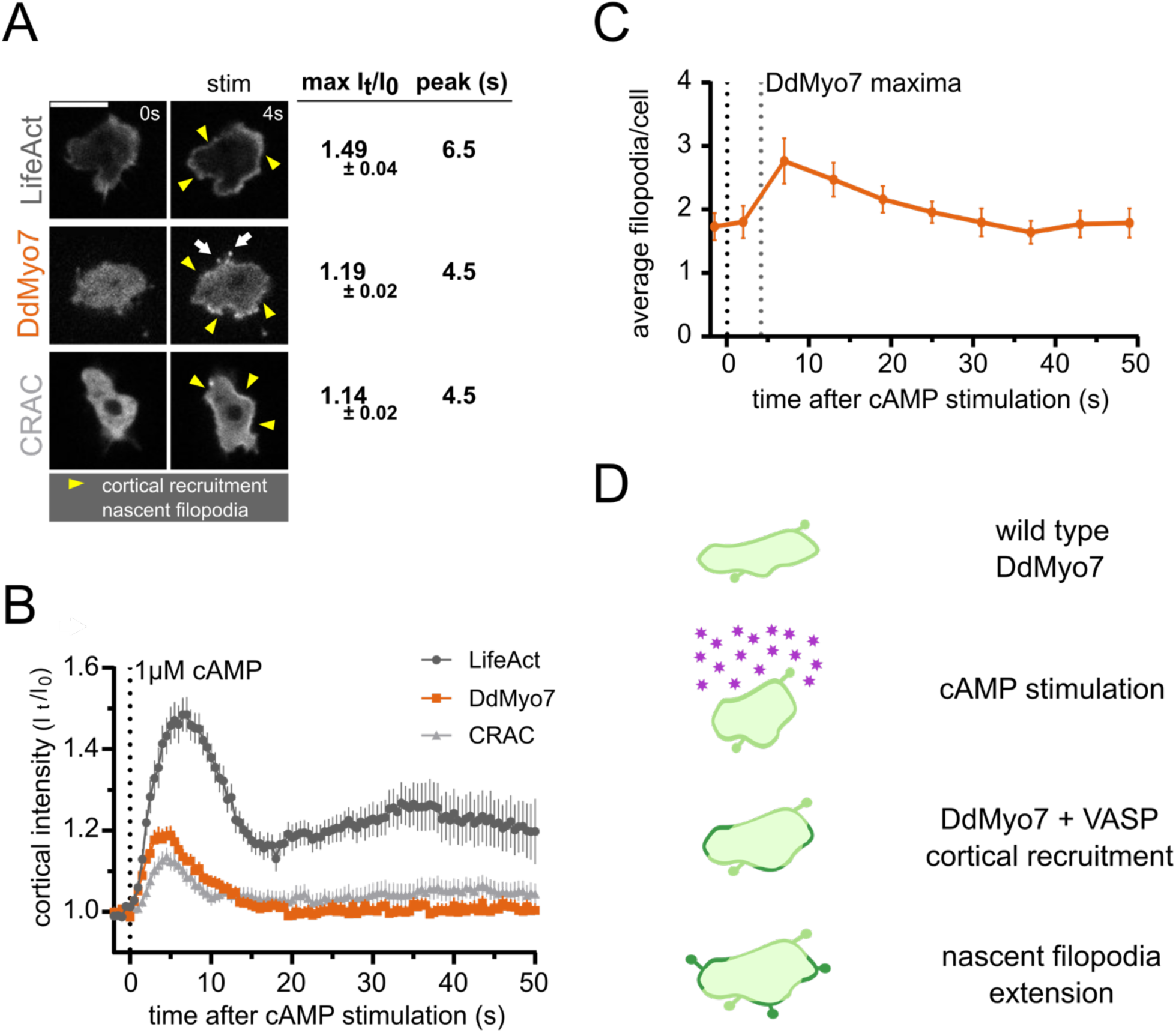
Global stimulation with chemoattractant cAMP results in cortical recruitment of DdMyo7 and stimulates filopodia formation. **A.** Representative images of wild type *D. discoideum*cells overexpressing actin marker RFP-LifeAct, GFP-DdMyo7, or GFP-CRAC immediately prior to (0s), and 4 seconds after global cAMP stimulation. Yellow arrows indicate regions of cortical enrichment and white arrows indicate nascent filopodia tipped with GFP-DdMyo7. Max cortex/cytoplasm intensities and peak times ± SEM are shown to the right. Scale bar = 10 µm. **B.** RFP-LifeAct (n = 38 cells, 3 experiments), GFP-DdMyo7 (n = 55 cells, 2 experiments), and GFP-CRAC (n = 32 cells, 6 experiments) are rapidly recruited to the cortex following global cAMP stimulation. The cortical intensity I_t_= (cortex - background) / (cytoplasm - background). I_t_was normalized to time 0s (I_0_), immediately before cAMP stimulation. Error bars = SEM for all cells. **C.** The number of GFP-DdMyo7 tipped filopodia per cell over time after global stimulation with cAMP. Vertical lines indicate stimulation time and peak time for GFP-DdMyo7 cortical recruitment from panels A and B. Error bars = SEM. **D.** Model of DdMyo7 and VASP recruitment following cAMP stimulation and consequent extension of DdMyo7 tipped filopodia (light green = pre-existing filopodia and cortical actin; dark green = stimulation-induced filopodia and increased cortical actin).

### Directionally biased filopodia formation during chemotactic migration

The formation of filopodia following global cAMP stimulation suggested that chemotactic stimuli can bias the site of filopodia production. While enrichment of actin at the leading edge of a polarized chemotactic *Dictyostelium* has been well-described (e.g. (Gerisch et al., 1995)), the location of filopodia in amoebae undergoing directed migration has yet not been reported. Control cells expressing GFP-DdMyo7 were placed in a well of the assay plate containing an established cAMP gradient and both cell migration and location of filopodia of cells moving under the agarose analyzed. The position of filopodia made by cells moving with high persistence in a stable gradient was translated on to an xy-plot (Fig 3B, Supp Fig 3; see Methods) with 0° being set to the direction of the source of the chemoattractant. A total of 2364 filopodia events were detected (74 cells, 6 independent experiments) and the vast majority of the observed filopodia events were biased towards the cAMP source (Fig. 3A&C). Dividing the angular coordinates of the cell periphery into quadrants reveals this enrichment toward the source of cAMP gradient is statistically significant (*X*^2^ (DF = 3, *N* = 2364 filopodia) = 366.5, *p*< 0.0001 (Fig. 3C). A peak number of these filopodia events (254, ∼11% of total) originate from a highly localized region of the perimeter oriented toward the gradient source (∼2.7% of the perimeter, 9.7°/360°; Fig 3C).

**Figure 3.**
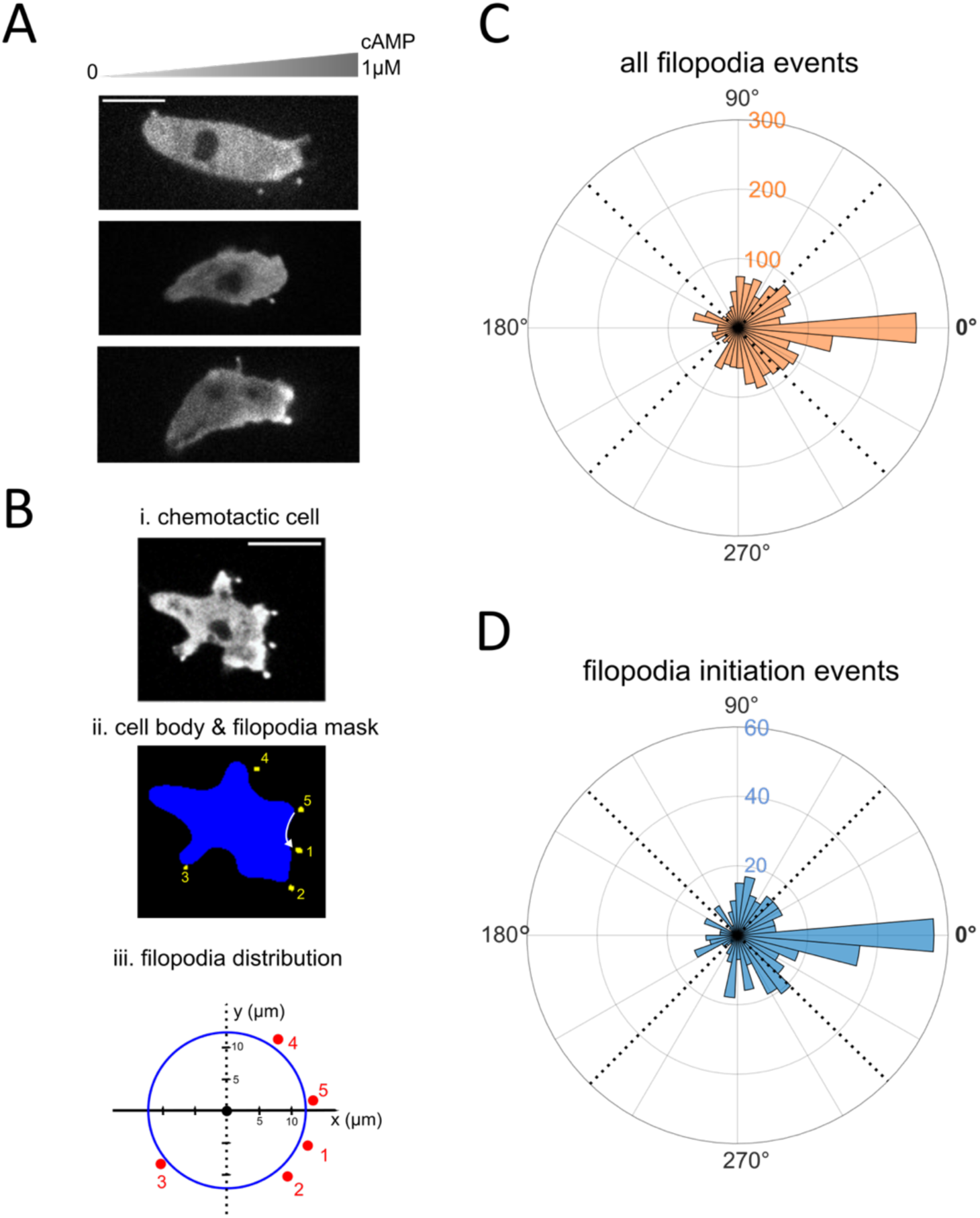
Filopodia formation is biased towards a chemotactic source. **A.** Representative micrographs of *myo7* null cells expressing GFP-DdMyo7 chemotaxing towards 1µM cAMP under agarose. Scale bar = 10 µm. **B.** Example of the arc length analysis. The closest pixel of the cell mask (*ii*, blue) to the tip mask center (*ii*,yellow) is assigned as the filopodia shaft. Arc length is measured from the right most pixel (*ii*, white arrow for tip 1). Arc lengths are then visualized as circumference fraction (*iii*). Scale bar = 10 µm. **C** and **D.** Arc length rose plot of all filopodia events where degrees are normalized arc lengths. **C.** Rose plot of all filopodia events detected during chemotaxis of *myo7* null cells expressing GFP-DdMyo7 (n = 74 cells, 6 experiments). The 0° position points towards higher cAMP concentrations, and the radial axis is the number of filopodia events in each bin (9.7° per bin). Dotted lines indicate the quadrant relative to the gradient. The relation between quadrant and total filopodia was significant, *X*^!^(DF = 3, *N* = 2364 filopodia) = 366.5, *p*< 0.0001. **D.** Subset of panel C, plotting only filopodia initiation arc lengths, where initiation events are defined as the first time point a filopodium appears. The relation between quadrant and initiation events was significant, *X*^!^(DF = 3, *N* = 458 initiation events) = 81.83, **** *p*< 0.0001.

The observed filopodia could have either arisen *de novo* or been selectively stabilized by the gradient so to address this, the data was parsed to examine filopodia initiation events. Out of a total of 458 initiation events the majority were in the leading quadrant, with a full 12%, of filopodia initiating in just the 3% perimeter arc length bin closest to the gradient (Fig 3D). These data along with the time lapse data reveal that cells most often from new filopodia at the leading edge. It also shows that some filopodia remained adhered to the substrate as cells continue to move up the gradient, with the adhering filopodia now flanking the sides of the cell before retracting.

The analysis of filopodia orientation during chemotactic migration revealed that *Dictyostelium* initiate and maintain filopodia in the direction of cAMP source. This striking bias of filopodia initiation and localization towards the source of the chemotactic gradient is similar to what has been seen in neural crest cells where filopodia orientation becomes progressively polarized as cells move up a gradient in vivo (Genuth et al., 2018). This suggests that filopodia formation up a gradient maybe be an evolutionarily conserved adaptation during chemotaxis and cell migration.

### Directional migration of amoebae does not require filopodia

The biased location of the filopodia of chemotaxing cells suggested that these structures may play a role as sensors of the cAMP gradient. Wild type (AX3) and cells without filopodia (*myo7* null and *vasp* null) were placed in a well in an under-agarose assay and the chemotactic index, persistence, speed and polarization (circularity) of each cell type were quantified. Despite the loss of filopodia, both the *myo7* and *vasp* null mutants migrated directionally with chemotactic indices similar to that of wild type cells (Fig. 4A, D), revealing that filopodia are not necessary for sensing cAMP (Fig. 4B-D). Consistent with the wild type chemotactic indices, the persistence of the *myo7* and *vasp* null mutants was comparable to wild type cells (Fig. 4D). Chemotactic cells are typically highly polarized and, interestingly, *myo7* and *vasp* null mutants were more polarized (less circular) than wild type Ax3 Figure 4A and Supp Fig 4D). This suggests that the dynamic extension of filopodia contributes to changes in cell shape (i.e. extending new pseudopodia) during chemotactic migration.

**Figure 4.**
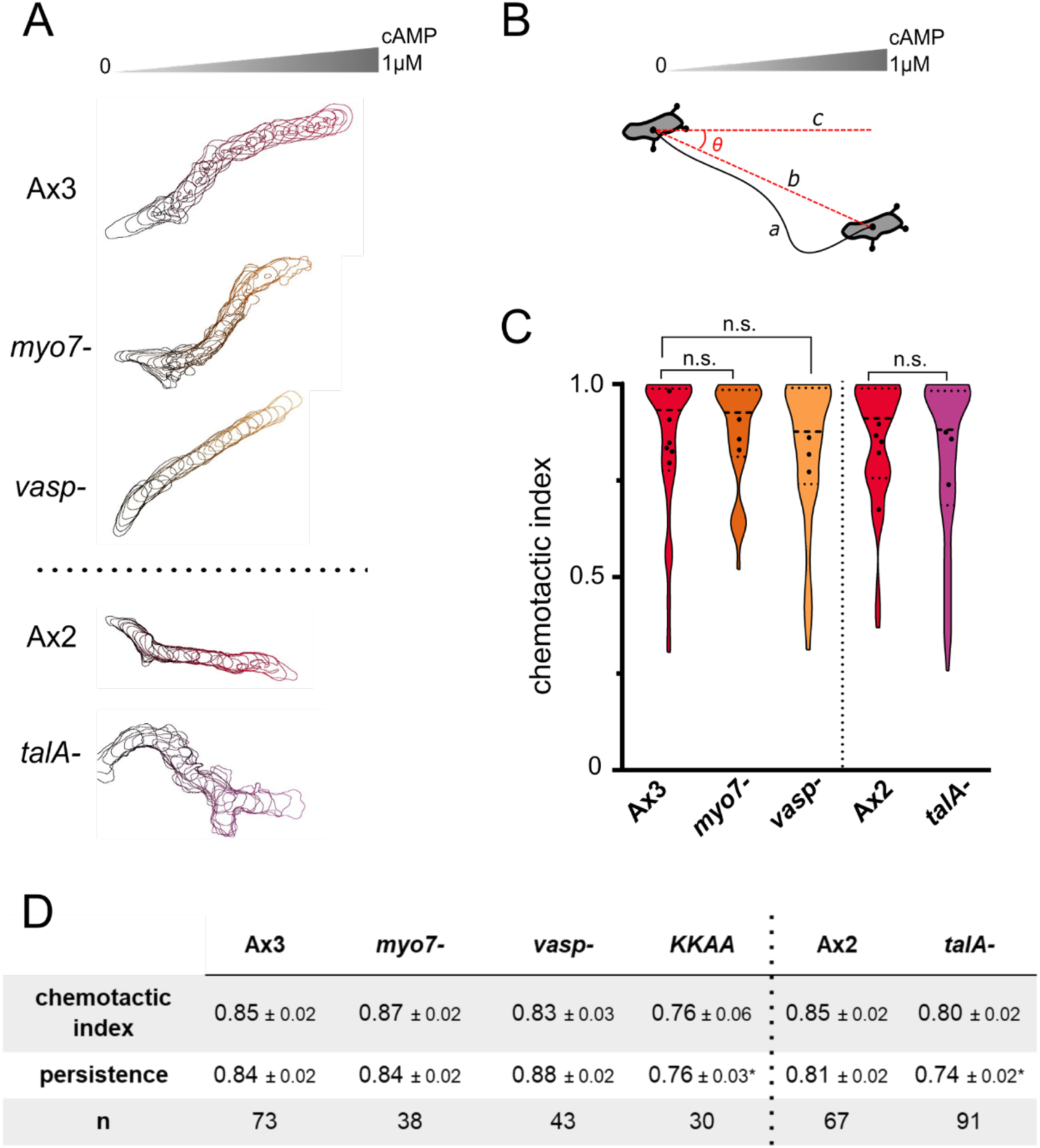
Filopodia are dispensable for directional and persistent chemotaxis. **A.** Representative perimeter plots for different genotypes migrating under agarose in a 1µM cAMP gradient for 5 minutes. **B.** Depiction of chemotaxis parameter calculations. Total distance traveled = line *a*. Displacement = line *b*. Chemotactic gradient vector = line *c*. Chemotactic index = cosine of the angle between *b* and *c*. Persistence = *b* / *a*. **C.** Average chemotactic index for each mutant. Dashed line separates genetic backgrounds Ax3 and Ax2 for the different mutant lines. Violin plots display kernel density distributions of individual cells; dashed line = median; dotted lines = quartiles. Black circles represent means of individual experiments. n.s. = no statistical significance (Ax3: Kruskal-Wallis ANOVA and multiple comparisons test, p > 0.99; Ax2: Mann-Whitney U test, p = 0.32). **D.** Summary of chemotaxis parameters. *n* = number of cells per genotype, combined from multiple individual experiments. One way ANOVA and multiple comparisons test (Ax3: * p < 0.05; **** p < 0.0001), and unpaired t test (Ax2; * p < 0.05).

### Migration and filopodia number

Chemotactic cells lacking filopodia migrate significantly slower than wild type Ax3 (Fig. 5C, Supp Figure 4D). Since the filopodia mutants *myo7* and *vasp* null mutants exhibit reduced substrate adhesion (Tuxworth et al., 2001, Han et al., 2002), the decreased speed could be simply attributed to weaker overall contact with the substrate. To test this, the *talA* null mutant was examined because it has an adhesion defect yet still produces filopodia (Niewöhner et al., 1997, Gebbie et al., 2004, Tuxworth et al., 2005). Chemotaxing *talA* cells have the same speed as wild type (Figure 5C, Supp Fig 4D). The only difference between the *talA* null and wild type cells migrating in a gradient is that they have slightly decreased persistence (Fig 4D). This may be because close adhesion at the front of the cell is not required for normal migration speeds. It is also possible that TalB, a TalA homologue upregulated during early development that is strongly localized to the leading edge of polarized cells (Tsujioka et al., 2008, Tsujioka et al., 2019), takes over its role during chemotactic migration.

**Figure 5.**
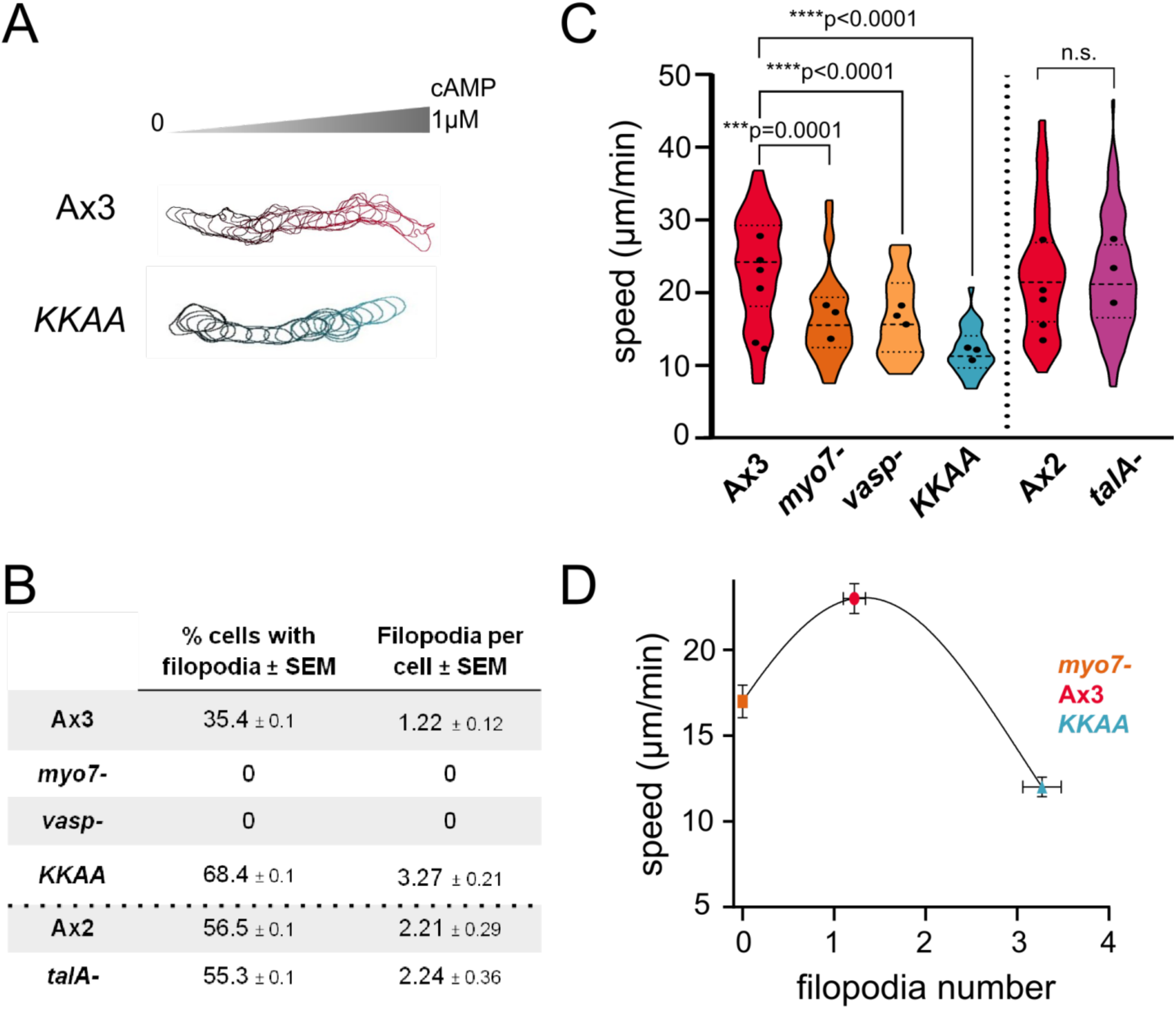
Filopodia number tunes the speed of chemotactic migration. **A.** Perimeter plots of Ax3 and DdMyo7 mutant KKAA migrating under agarose in a 1µM cAMP gradient over 5 minutes (n=30, 3 experiments). **B.** Average filopodia made per chemotaxing cell. **C.** Speed = total distance traveled (see Fig 3B, line *a*) / total time. Kruskal-Wallis ANOVA and multiple comparisons test (Ax3: *** p = 0.0001; **** p < 0.0001), and unpaired t test (Ax2; n.s. p = 0.76). Dashed line separates genetic backgrounds Ax3 and Ax2 for the different mutant lines. **D.** Average speed of all cells of a given genotype: *myo7-*, Ax3, *KKAA* plotted as a function of filopodia number in filopodia-positive cell subset. The line drawn on the plot is to illustrate the trend of the data.

The reduced speed of the mutants lacking filopodia suggested that increasing their number might enhance their motility. DdMyo7 is regulated by head-tail autoinhibited and mutation of key conserved residues in the tail (KKAA) shifts the equilibrium towards the open, active state resulting in an a 2-fold increase in filopodia in randomly moving cells (Petersen et al., 2016, Arthur et al., 2019). Chemotaxing KKAA DdMyo7 cells also produce increased numbers of filopodia (Fig. 5B). Unexpectedly, the KKAA DdMyo7 cells moving in a cAMP gradient exhibited a significant decrease in migration speed compared to wild type Ax3 (Fig. 5C, Supp. Fig. 4D). The average speed of KKAA DdMyo7 was also significantly slower than that either the *myo7* or *vasp* nulls (Fig 5C, Supp Fig 4D). This revealed that a nonlinear relationship exists between chemotactic speed and filopodia number (Fig. 5D, Supp Fig S3). Specifically, the fastest and presumably optimized, migration rates occur at wild type filopodia numbers. These data suggest that filopodia number have e migration.

## Conclusion

Fast-moving chemotaxing amoebae do not require filopodia for persistent, directional migration in a gradient. Instead, filopodia number impact cell migration speed, with changes in numbers affecting efficient migration in a gradient. Migration relies on the coordinated extension of a pseudopod towards the gradient source followed by rear retraction (Ridley et al., 2003). The pseudopod must make stable contact with the substrate in order for the cell the move forward. The biased extension of filopodia from the leading edge, towards the gradient source during chemotaxis, suggests that they aid in keeping the front of the cell in contact with the substrate, promoting optimal movement. The migration speed of amoeboid cells does not scale linearly with filopodia number (Fig 5D) and there instead appears to be an optimal number of filopodia needed for the most efficient movement. Interestingly, this is reminiscent of the relationship between integrin-mediated adhesion and cell migration speeds in slower moving mesenchymal cells (Palecek et al., 1997). It suggests that role of filopodia in amoeboid migration could be to mediate optimal contact between the leading edge of the cell both during random motility and chemotactic migration when cells are either hunting for food or aggregating together during the early stages of multicellular development.

## Acknowledgements

We thank Ashley Arthur and Casey Eddington for many helpful discussions and Ashley Arthur for feedback on the manuscript. This work was supported by a University of Minnesota Interdisciplinary Doctoral Fellowship to YS and a grant from the NIH National Institute of General Medical Sciences (GM122917) to MAT

## Author contributions

Conceptualization, MAT; Methodology, YS, LDS; Investigation, LDS, ALS, YS; Writing – Original Draft, ALS, LDS; Writing – Review & Editing, ALS, LDS, YS, MAT; Visualization, ALS, LDS, YS; Software LDS, YS; Validation YS, LDS; Formal Analysis ALS, LDS, YS; Funding Acquisition, MAT; Resources, ALS, LDS; Supervision, MAT; Project Administration MAT

## Materials & Methods

*Dictyostelium discoideum* cells (Ax2 and Ax3 background) were grown at 22°C on bacteriological plastic plates in HL5 glucose medium (Formedium) supplemented with 10 kU/mL penicillin G and 10 µg/mL streptomycin sulfate (Sussman, 1987). The Ax3 non-homologous control and *myo7* null lines, as well as the *talA* and *vasp* null cells have been described previously (Niewöhner et al., 1997, Tuxworth et al., 2001, Han et al., 2002). Cells were transformed with exogenous expression plasmids of interest by electroporation (Gaudet et al., 2007), and were selected for and maintained using 10 µg/mL G418 (ThermoFisher) and/or 35 µg/mL hygromycin B (Gold Bio). Plasmids and cell lines are listed in Supp Table 1.

Chemotactically competent cells were prepared in one of two ways. Method 1: cells were collected and washed in development buffer (5mM Na_2_HPO_4_, 5mM KH_2_PO_4_, 1mM CaCl_2_, 2mM MgCl_2_, pH 6.4) and resuspended at a concentration of 1-2 × 10^7^ cells/mL. Cells were shaken at 150 RPM at 22°C and pulsed with 100nM cAMP every 6 minutes after the first hour of starvation (Parent et al., 1998). After shaking for 4 hours, cells were collected, washed twice in PB, and diluted to 1 × 10^6^ cells/mL before being added to the outer wells of the under-agarose imaging dish. Method 2: cells were collected and washed in development buffer and 2 × 10^6^cells were plated in a 35mm plastic bacteriological plate. Plates were placed at 12°C for 8 - 14 hours, then moved to room temperature for an hour (Brzeska et al., 2012) before being resuspended and added into the outer wells of the under-agarose imaging dish.

### Under-agarose migration assays

Under-agarose chemotaxis experiments were performed as previously described (Laevsky and Knecht, 2001, Brazill, 2016). Briefly, 0.7 - 0.8% LE agarose (Genemate, Bioexpress) was dissolved into PB and solidified for 1 hr in a 35 mm glass-bottom imaging dish (Mattek). Three wells (approximately 2mm × 10mm) were cut into the agarose above the glass bottom with a razor. For chemotaxis assays, 1 µM cAMP was loaded into the middle well (approximately 75 µl) for 60 min to establish the gradient.

### cAMP global stimulation experiments

Log-phase cells were collected and washed into LPS buffer (40 mM Na_2_PO_4_, 20 mM KCl, 0.24 mM MgCl_2_, 0.34 mM streptomycin sulfate, pH 6.4). Cells were plated at a density of 3.3 - 3.6×10^6^ cells/cm^2^ on a stack of damp filters consisting of two Whatman No. 17 absorbent pads topped with one black Whatman No. 29 filter paper and maintained at 22°C in a humidity chamber. Cells were scraped into phosphate buffer (PB; 16.8 mM sodium/potassium phosphate, pH 6.4) after five hours and washed twice into PB prior to live-cell imaging (Sussman, 1987). Cells were adhered to 35 mm glass imaging dishes at a density of 0.5 - 1 × 10^5^ cells/cm^2^ in PB for 10 minutes. Cells were imaged at 2 fps for 10 sec prior to and 50 sec post global stimulation (final concentration of 1µM cAMP). Fluorescein sodium salt or Alexa Fluor 594 Hydrazide (Sigma) was added to cAMP stock for stimulation frame detection (opposite channel to fluor-tagged protein of interest).

*Microscopy*. Confocal microscopy was performed with 40x and 63x Ph3 Plan-Apochromat oil-immersion objectives (NA 1.4) on a Marianas Spinning Disk Confocal imaging system (3i) based on a Zeiss Axiovert microscope. This system was controlled by SlideBook 6.0 software (Intelligent Imaging Innovations) and equipped with a confocal scanner unit (Yokogawa CSU-X1), an electron-multiplying CCD camera for super-resolution (Photometrics Evolve 512), a monochrome CCD camera for quantitative fluorescence microscopy (Photometrics CoolSNAP HQ2), a motorized stage controller (ASI MS-2000), and laser lines at 488 and 561 nm.

For chemotaxis migration assays, all cells were imaged approx. 1 hour after being added to dish; one frame was captured every 15 sec for 15min. For cells used to analyze filopodia formation in a gradient, cells were imaged at a rate of 2 frames per sec for 2 min, with 2 z planes 0.5 µm apart. Under agarose experiments for cell lines not expressing fluorescent proteins were imaged with 40x oil immersion objective (NA 0.55) on a Zeiss Axiovert microscope.

### Image processing and analysis

Confocal images were analyzed using the custom Java-based FIJI/ImageJ macro Seven (Schindelin et al., 2012, Petersen et al., 2016). This suite of programs and macros was updated to calculate the cortex/cell intensity ratio (It) over time after cAMP stimulation. Cell masks were generated for every frame by thresholding in ImageJ, and the grayscale values of the cytoplasm, cortex, and background were measured (arbitrary units). The background intensity was subtracted from the raw cortex and cytoplasm intensities to correct for photobleaching and for each time point, It = (cortex - background) / (cytoplasm - background). The output from Seven was processed in RStudio and compiled as replicates in Graphpad Prism.

Filopodia formation in a gradient was quantified using a set of custom MATLAB scripts, collectively called Space7. Random Forest classifier (Ilastik 1.3.3; (Berg et al., 2019)) was trained with three classes: background, cell body, GFP-DdMyo7 puncta. All images were then run through the classifier in the batch mode. Cell bodies object classification was performed by hysteresis method with a smoothing parameter 5.0 and threshold core parameter of 0.80 and final of 0.5. Puncta object classification was done with the threshold parameter 0.4 and smoothing parameter 1. Cell body object predictions were imported to FIJI/ImageJ macro to create coordinates of the cell outline. Cell outline coordinates and puncta centroids were then used to find the filopodia shaft pixel on the cell outline (closest pixel on the outline to the puncta centroid). Filopodia longer than 3 µm were discarded to avoid over counting. Arc length in µm was calculated starting from the rightmost pixel clockwise. In order to analyze the arc length dynamics over time, arc length was normalized to cell perimeter (Fig. 3B (iii), perimeter of the blue circle corresponds to perimeter of the cell in Fig. 3B (i)) in degrees.

Under agarose chemotaxis was tracked using a combination of Ilastik 1.3.3 and a custom FIJI/ImageJ macro, that identified cell edges and masked DIC images over time (Schindelin et al., 2012, Berg et al., 2019). The x/y values for each cell were processed in RStudio. All parameters are based on the translocation of the centroid of the cell. Chemotactic index was calculated as final distance travelled towards the cAMP (x component only) divided by displacement (Chen et al., 2003, Brazill, 2016). Persistence was calculated as displacement divided by total distance travelled. Speed was calculated as total distance travelled divided by time. Data was plotted using Graphpad Prism and imported into InkScape for vector processing and design.

Cells were excluded from analysis if they were in frame for less than 2min or counted for more than 15min (max time). Because the gradient generated using under agarose experiments is dissipating over time and cells act as competing sources of cAMP production, only cells with positive chemotactic index were analyzed (additionally, all definite outliers were removed, Rout test; Q=0.1%). Cells traveling at a speed less than 5 µm/min were excluded, as that is the average speed of vegetative cells (Fig 1).

### Quantification and statistical analysis

All groups were tested for normality of distribution (D’Agostino and Pearson test). All tests were done to calculate significance at the alpha = 0.05 level. When comparing two genotypes across one parameter, either T test or Mann Whitney U test were used; One way ANOVA or Kruskall Wallis and respective multiple comparison tests were used when comparing one parameter across more than 2 genotypes.

## SUPPLEMENTARY FIGURES

**Supplementary Figure S1.**
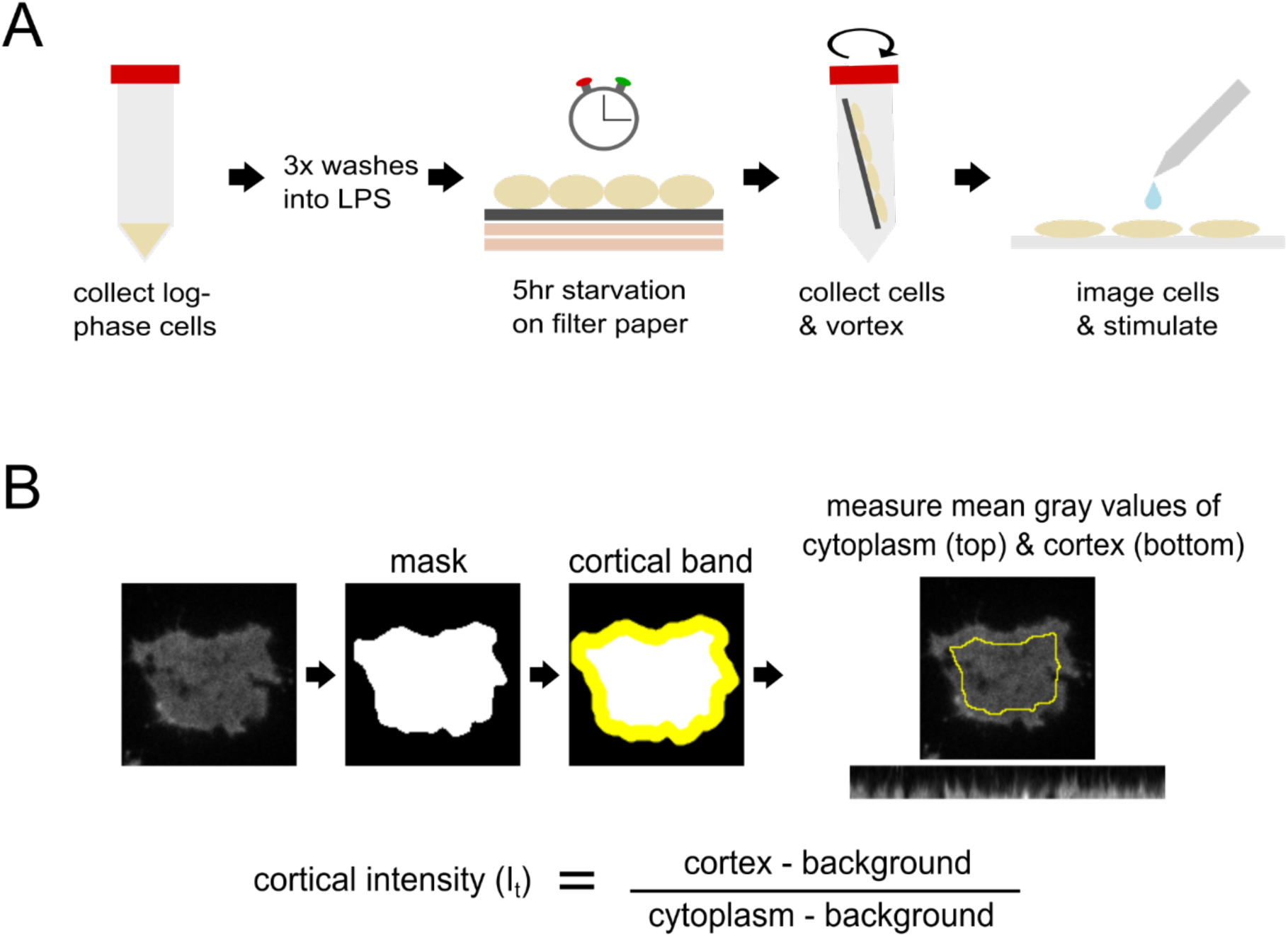
cAMP stimulation experimental set up and analysis pipeline. **A.** Cartoon of cAMP stimulation experimental protocol. **B.** Pipeline for cortical intensity calculation for cAMP stimulation experiments.

**Supplementary Figure S2.**
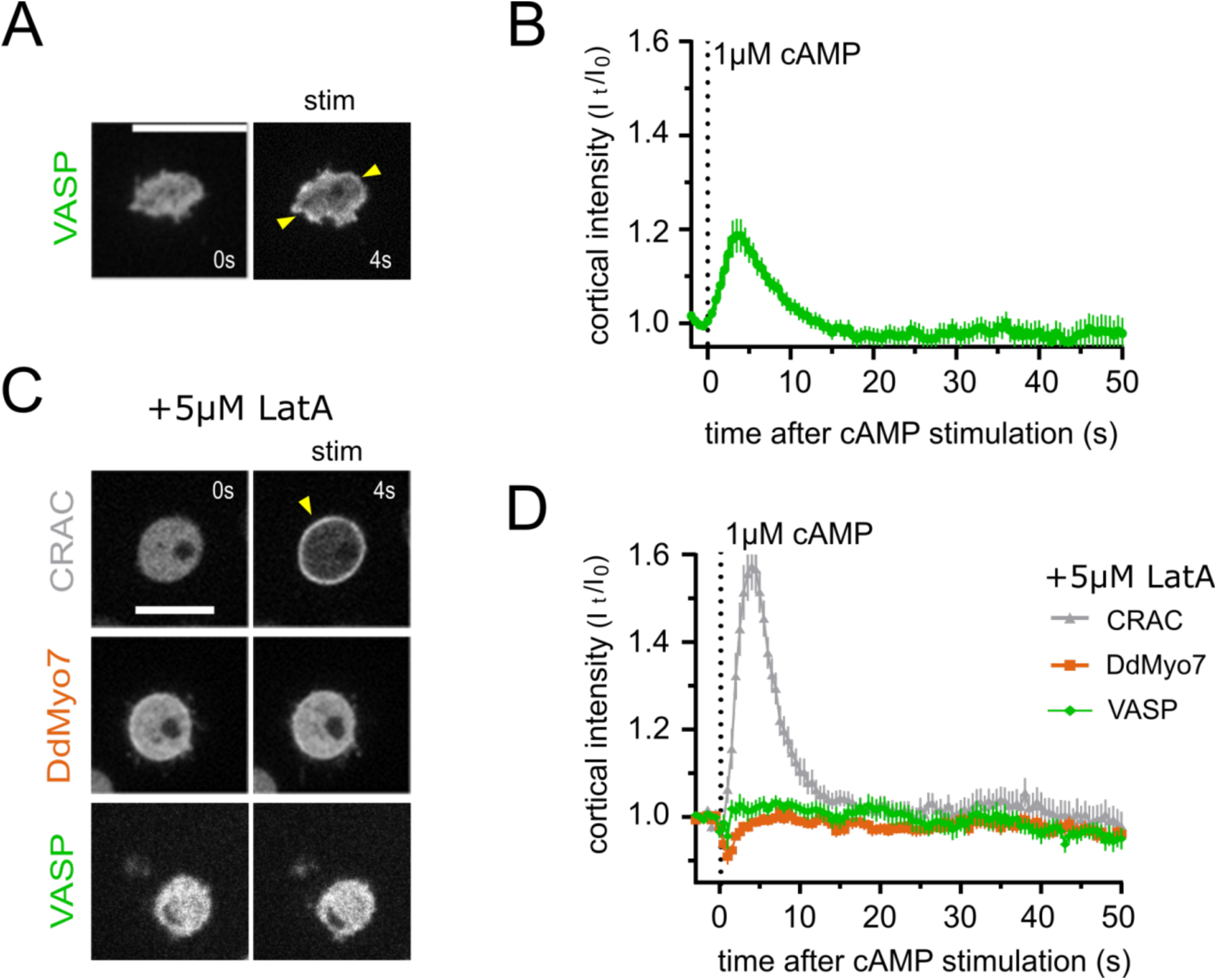
DdMyo7 and VASP cortical recruitment is actin dependent. **A.** Representative image of wild type cells expressing mApple-VASP prior to and 4 seconds post global cAMP stimulation. Yellow arrows indicate regions of cortical enrichment. Scale bar = 10 µm**. B.** Quantification of transient cortical recruitment of VASP following stimulation (n = 37 cells, 6 experiments). Error bars = SEM for all cells. **C.** Representative micrographs of wild type cells expressing GFP-CRAC, GFP-DdMyo7, or mApple-VASP after treatment with 5µM LatA immediately prior and 4 seconds post global cAMP stimulation. Yellow arrow indicates regions of cortical enrichment. Scale bar = 10 µm. **D.** Quantification of transient cortical recruitment of CRAC, DdMyo7, and VASP following stimulation (GFP-CRAC: n = 25 cells, 6 experiments; GFP-DdMyo7: n = 29, 4 experiments; mApple-VASP: n=25, 5 experiments). Error bars = SEM for all cells.

**Supplementary Figure S3.**
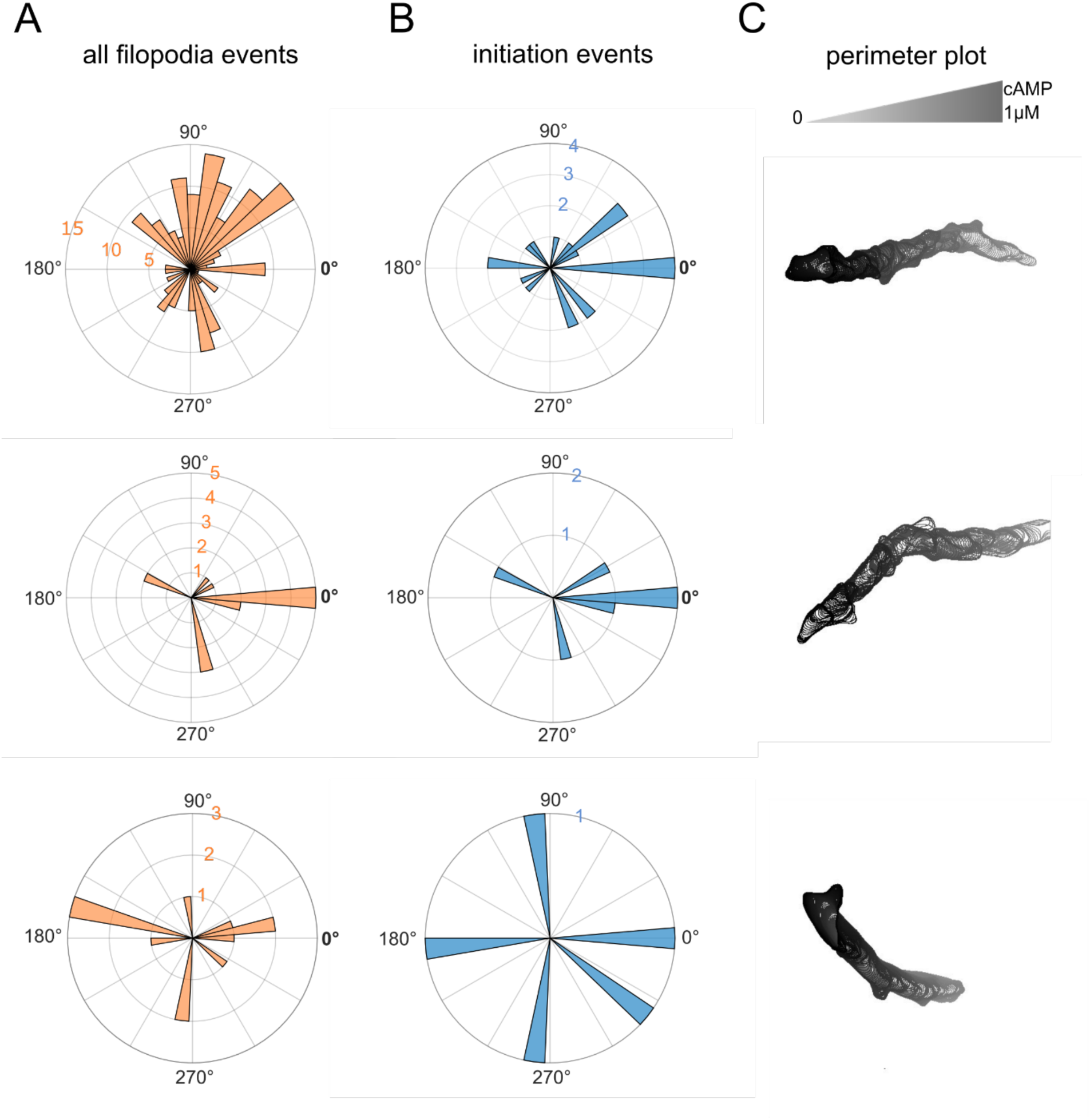
Representative Analysis of individual cells migrating in a cAMP gradient. Shown are their respective (A) all filopodia events (B) initiation events and (C) perimeter plots. 0° on rose plots point towards higher cAMP concentrations, and the radius of each circle corresponds to the number of filopodia/initiation events in each bin (9.7° per bin).

**Supplementary Figure S4.**
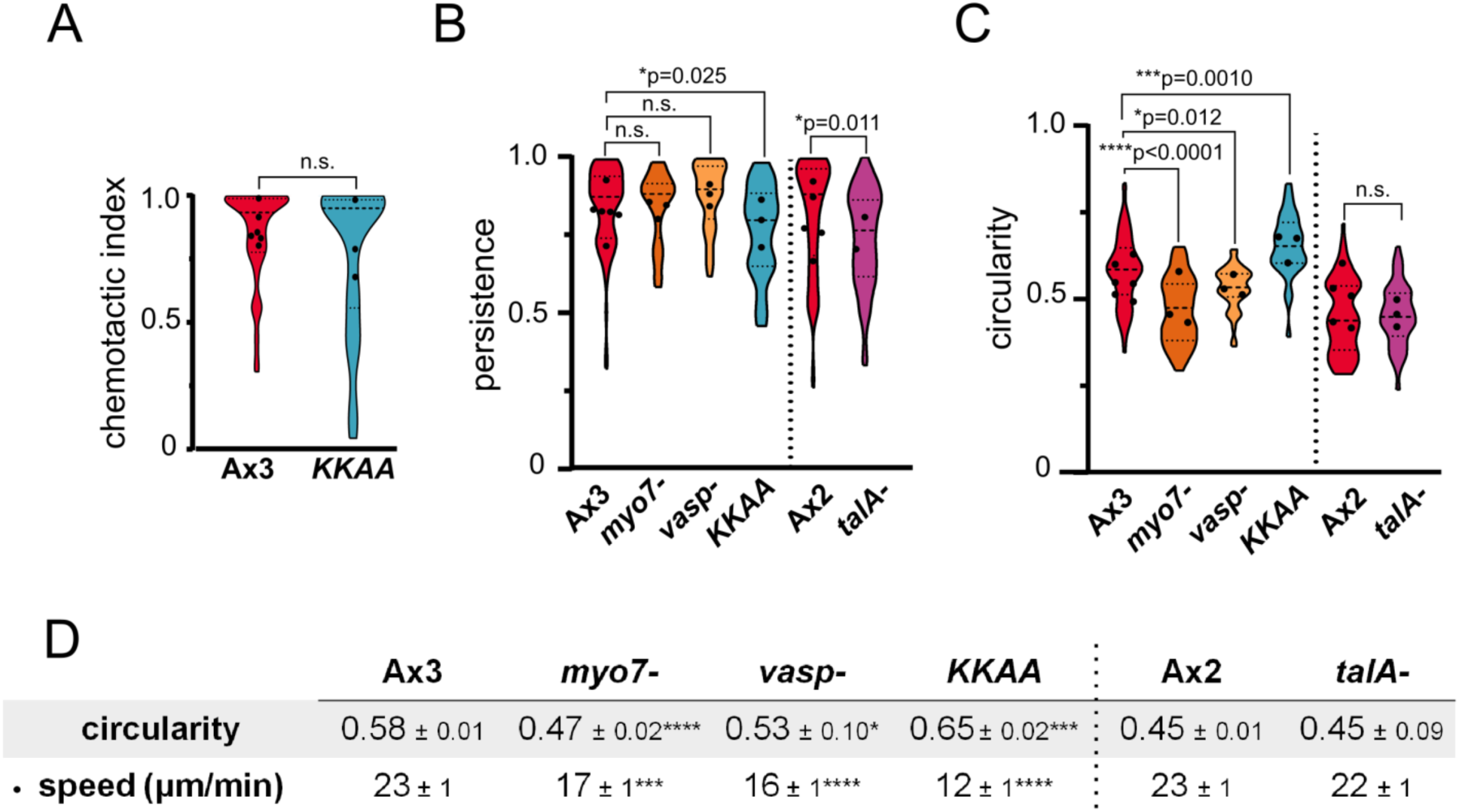
Quantification of the chemotactic index, persistence and circularity of cells migrating under agarose in a cAMP gradient. **A.** Average chemotactic index of Ax3 and *KKAA* mutant. All violin plots in figure display kernel density distributions of individual cells; dashed line = median; dotted lines = quartiles. Black circles represent means of individual experiments. n.s. = no statistical significance (Kruskal-Wallis ANOVA and multiple comparisons test, p > 0.99). **B**. Average persistence for each mutant. Dashed line separates genetic backgrounds Ax3 and Ax2.One way ANOVA and multiple comparisons test (Ax3: n.s. p > 0.05; * p < 0.05), and unpaired t test (Ax2; * p < 0.05). **C.** Average circularity for each mutant. Circularity = 4π (area /perimeter^2^). Value of 1 indicates circle, those closer to 0.5 correspond to an ellipse.One way ANOVA and multiple comparisons test (Ax3: * p < 0.05; *** p < 0.005; **** p < 0.0001), and unpaired t test (Ax2; n.s. p = 0.89). **D.** Quantified mean circularity and speed for each genotype.

**Supplemental Table 1.**
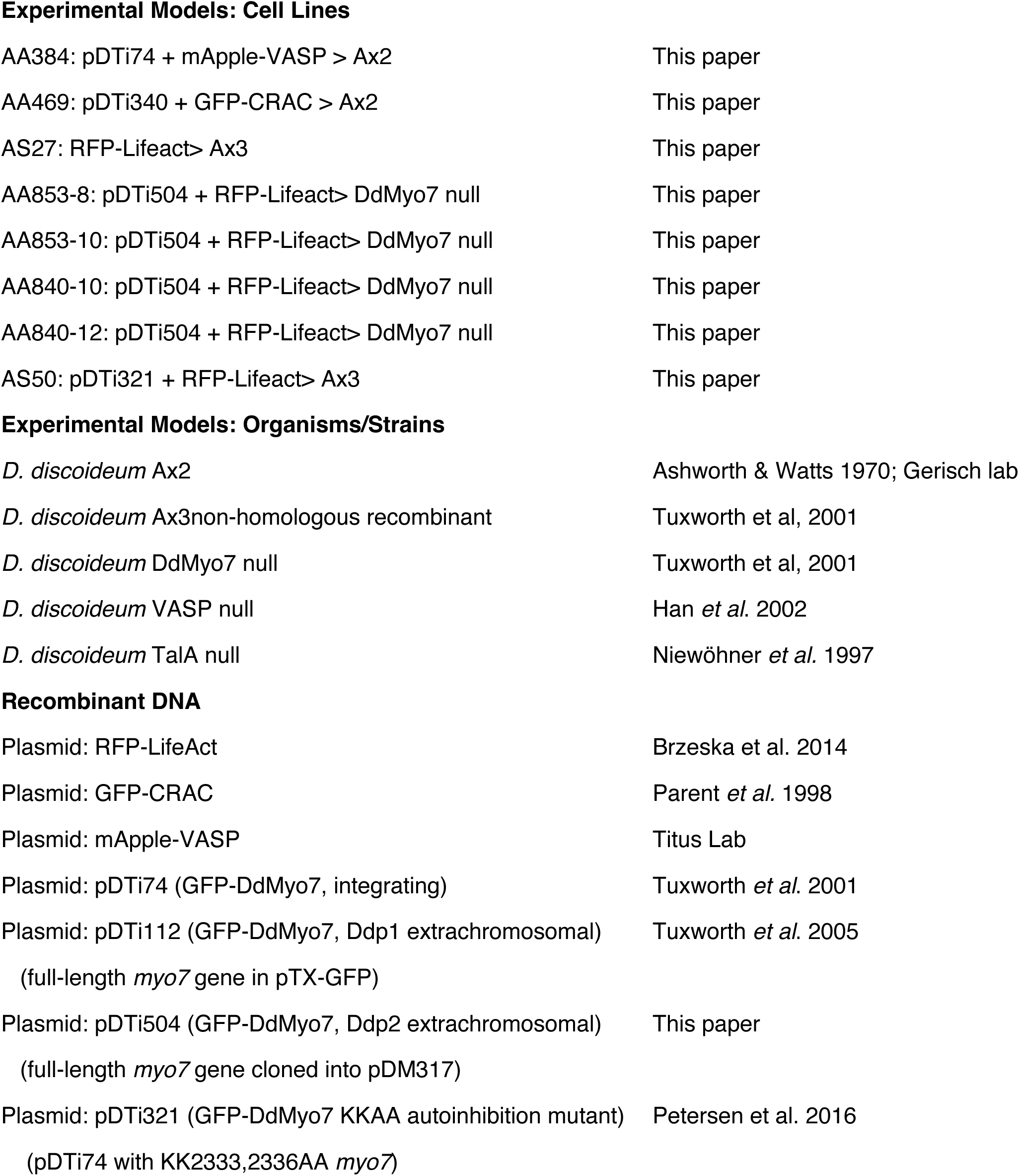

Experimental Models: Cell Lines
|  |  |
| --- | --- |
| AA384: pDTi74 + mApple-VASP > Ax2 | This paper |
| AA469: pDTi340 + GFP-CRAC > Ax2 | This paper |
| AS27: RFP-Lifeact> Ax3 | This paper |
| AA853-8: pDTi504 + RFP-Lifeact> DdMyo7 null | This paper |
| AA853-10: pDTi504 + RFP-Lifeact> DdMyo7 null | This paper |
| AA840-10: pDTi504 + RFP-Lifeact> DdMyo7 null | This paper |
| AA840-12: pDTi504 + RFP-Lifeact> DdMyo7 null | This paper |
| AS50: pDTi321 + RFP-Lifeact> Ax3 | This paper |
| Experimental Models: Organisms/Strains |  |
| *D. discoideum* Ax2 | Ashworth & Watts 1970; Gerisch lab |
| *D. discoideum* Ax3non-homologous recombinant | Tuxworth et al, 2001 |
| *D. discoideum* DdMyo7 null | Tuxworth et al, 2001 |
| *D. discoideum* VASP null | Han *et al.* 2002 |
| *D. discoideum* TalA null | Niewöhner *et al.* 1997 |
| Recombinant DNA |  |
| Plasmid: RFP-LifeAct | Brzeska et al. 2014 |
| Plasmid: GFP-CRAC | Parent *et al.* 1998 |
| Plasmid: mApple-VASP | Titus Lab |
| Plasmid: pDTi74 (GFP-DdMyo7, integrating) | Tuxworth *et al.* 2001 |
| Plasmid: pDTi112 (GFP-DdMyo7, Ddp1 extrachromosomal)<br>(full-length *myo7* gene in pTX-GFP) | Tuxworth *et al.* 2005 |
| Plasmid: pDTi504 (GFP-DdMyo7, Ddp2 extrachromosomal)<br>(full-length *myo7* gene cloned into pDM317) | This paper |
| Plasmid: pDTi321 (GFP-DdMyo7 KKAA autoinhibition mutant)<br>(pDTi74 with KK2333,2336AA *myo7*) | Petersen et al. 2016 |

## Notes

### Competing Interest Statement

The authors have declared no competing interest.

